# Efficacy of a novel iodine complex solution, CupriDyne, in inactivating SARS-CoV-2

**DOI:** 10.1101/2020.05.08.082701

**Authors:** Emily Mantlo, Alex Evans, Laura Patterson-Fortin, Jenny Boutros, Richard Smith, Slobodan Paessler

## Abstract

The coronavirus known as SARS-CoV-2, which causes COVID-19 disease, is presently responsible for a global pandemic wherein more than 3.5 million people have been infected and more than 250,000 killed to-date. There is currently no vaccine for COVID-19, leaving governments and public health agencies with little defense against the virus aside from advising or enforcing best practices for virus transmission prevention, which include hand-washing, physical distancing, use of face covers, and use of effective disinfectants. In this study, a novel iodine complex called CupriDyne^®^ was assessed for its ability to inactivate SARS-CoV-2. CupriDyne was shown to be effective in inactivating the virus in a time-dependent manner, reducing virus titers by 99% (2 logs) after 30 minutes, and reducing virus titers to below the detection limit after 60 minutes. The novel iodine complex tested herein offers a safe and gentle alternative to conventional disinfectants for use on indoor and outdoor surfaces.

## Introduction

Beginning in December of 2019, the novel coronavirus known as Severe Acute Respiratory Syndrome Coronavirus 2 (SARS-CoV-2) has caused a global pandemic wherein over 3.5 million people have been infected, and over 250,000 people have died of the disease known as COVID-19 as of May, 2020 (Johns Hopkins Coronavirus Resource Center, 2020). First identified in patients in Wuhan City in Hubei province, China (Lu *et al*, 2020), COVID-19 causes symptoms including a dry cough, fever, shortness of breath and sore throat (CDC, 2020), and in severe cases can cause severe pneumonia, pulmonary edema, organ failure, and death (Chen *et al,* 2020). The pandemic has spread to over 180 countries, with most governments enacting unprecedented restrictions on movement and assembly of people to curb the growth of the pandemic. Currently, there is no vaccine or effective treatment for COVID-19.

Public health organizations have provided guidance on best practices to limit the spread of COVID-19, which include frequent hand washing, physical distancing, the use of face covers, and the use of effective cleaners and disinfectants to decontaminate surfaces (CDC, 2020; WHO, 2020). In the United States, the Environmental Protection Agency (EPA) has created a list of registered disinfectants that meet its criteria for efficacy against SARS-CoV-2 (US EPA, 2020).

Broadly, the most common disinfectant used against the SARS-CoV-2 is diluted bleach, a widely available and highly effective viral disinfectant. Bleach has several downsides when considered for widespread use including common irritation of skin, mucosal membranes, and airways, its instability when exposed to light, and its high degree of reactivity with organic material (WHO, 2014). Other common disinfectants such as ethanol and isopropyl alcohol are also effective and widely available, but also have downsides such as flammability, rate of evaporation, and skin sensitivity (Stanford, 2020). In light of these disadvantages, there is a need to develop new disinfectants that are effective against SARS-CoV-2 but which do not cause sensitivity or irritation issues among exposed people, and which can provide long-lasting efficacy.

Iodine-containing disinfectants have been used since the mid-19^th^ century (Sneader, 2005), and today iodine complexes such as povidone-iodine (PVP-I) are used in diverse applications that include surgical antiseptic, skin disinfection, and water disinfection (Durani and Leaper, 2008). While traditional Lugol’s iodine or iodine tinctures (wherein elemental iodine is dissolved in water or alcohol, respectively) are effective yet cytotoxic disinfectants, povidone-iodine complexes provide effective disinfection without significant cytotoxicity due to the slow release of iodine from the polyvinylpyrrolidone with which iodine is complexed. However, povidone-iodine causes noticeable staining of skin and surfaces and commonly causes skin irritancy (WHO, 2008), making it an unpopular hard surface disinfectant. Mechanistically, iodine is believed to react with and inactivate bacteria and viruses by oxidizing and/or iodinating critical proteins, DNA, RNA, and fatty acids (Gottardi, 1999; Gottardi, 2014). Iodine-containing solutions have been proven effective against a wide variety of viruses including influenza A, poliovirus, adenovirus type 3, mumps, and HIV (Wada *et al*, 2016; Kawana *et al*, 1997), with some indication of greater viricidal spectrum of activity compared to other commercially available disinfectants.

Iodine-containing solutions have historically been employed against other respiratory virus infection outbreaks. In 2006, PVP-I was used to inactivate the SARS coronavirus of the so-called SARS epidemic to below detectable levels in a laboratory study (Kariwa *et al*, 2006), and it was demonstrated effective against the Middle East Respiratory Syndrome Coronavirus (MERS-CoV) in 2015 (Eggers *et al*, 2015). Limited evidence also suggests iodine was successfully used to combat the spread of the 1918 H1N1 flu pandemic, also known as the Spanish Flu (Derry *et al*. 2009).

CupriDyne^®^ iodine complex (“CupriDyne”), made by Odor-No-More, Inc., a subsidiary of California-based life sciences company BioLargo, Inc., is a novel iodine complex solution that produces high local concentrations of iodine without causing the safety and staining issues associated with Lugol’s iodine or PVP-I respectively. CupriDyne uses a proprietary chemical solution to produce aqueous elemental iodine (I_2_) and cuprous iodide (CuI) in equilibrium.

CupriDyne is thought to react with airborne and surface-bound contaminants, microbes, and viruses through 1) direct oxidation and/or iodination by I_2_, and 2) reaction with cuprous iodide (CuI). In addition to the established antimicrobial action of I_2_, copper-containing complexes have been documented to exhibit bactericidal and viricidal activity (Borkow *et al*, 2004), providing a theoretical basis for added viricidal action of CupriDyne relative to traditional iodine solutions. In this study, CupriDyne was assessed for its efficacy in inactivating SARS-CoV-2 using a Vero cell monolayer infection model.

## Materials and Methods

### Cell lines and cell growth

Vero cells were grown in DMEM containing 1x L-glutamine, 1x penicillin/streptomycin, 1% MEM vitamins, and 10% FBS, and were incubated at 37° C in 5% CO_2_. Cells were used at 85-95% confluent growth.

### Virus stocks

The SARS-CoV-2 (USA-WA1/2020) virus was obtained from The World Reference Center for Emerging Viruses and Arboviruses (WRCEVA), University of Texas Medical Branch, Galveston, TX). SARS-CoV-2 was propagated in DMEM containing 1x L-Glutamine, 1x penicillin/streptomycin, 1% MEM vitamins, and 5% FBS at 37° C in 5% CO_2_. Prior to incubation, virus stocks were stored at −80° C at a concentration of 1×10^6^ TCID_50_ per mL. All experiments involving infectious virus were conducted by Dr. Slobodan Paessler’s laboratory at the University of Texas Medical Branch (Galveston, TX) in approved biosafety level 3 laboratories in accordance with institutional health and safety guidelines and federal regulations.

### Preparation of SARS-CoV-2 virus stocks

Briefly, SARS-CoV-2 was added at a MOI of 0.01 to Vero cells and incubated 1 hour at 37°C. Viral inoculum was removed, and fresh infection media added. Cells were incubated for an additional 48 hours before supernatant was collected. Supernatant was centrifuged for 5 minutes at 3000 rpm to remove cell debris. Virus stocks were stored at −80° C at a concentration of 1×10^6^ TCID_50_ per mL.

### Determination of Viral Titers

Viral titers were measured by TCID50 on Vero Cells. 96-well plates with Vero cells were used for this assay. Each log_10_ dilution of the virus was inoculated in quadruplicates. On day 4 post-infection, the cells were fixed with 10% formalin for 45 minutes and subsequently stained with crystal violet.

### Determination of virus inactivation using CupriDyne or controls

Aliquots of stock virus were mixed 1:10 by volume with CupriDyne iodine complex (CupriDyne (250 ppm, 25 ppm, or 2.5 ppm)) or control solutions, negative control (room temperature water) or positive control (boiling water >100° C)). The mixtures were incubated for predetermined time intervals at room temperature at which time infection media was added to neutralize antiviral activity. Subsequently, The SARS-CoV-2 viral titer (TCID_50_/mL) for each test substance was determined. The experiment was conducted in triplicate.

## Results

The ability of the CupriDyne iodine complex to inactivate SARS-CoV-2 was assessed using a Vero cell monolayer model. Undiluted (250 ppm), 1:10 diluted (25 ppm), and 1:100 diluted (2.5 ppm) solutions of CupriDyne were assayed in order to approximate the minimum concentration of iodine complex required to yield viricidal activity. The efficacy with which the CupriDyne iodine complex inactivated SARS-CoV-2 was shown to be both time- and concentration-dependent. Diluted samples of CupriDyne (25 ppm or 2.5 ppm) were not found to cause a statistically significant difference in SARS-CoV-2 titers. Undiluted CupriDyne (250 ppm) was shown to effectively inactivate the virus to a statistically significant extent after 10, 30, and 60 minutes.

After incubation with undiluted (250 ppm) CupriDyne for 10 minutes, viral titers dropped by 1 log (one-tailed t-test p-value = 0.0306). Viral titers dropped 2 logs (one-tailed t-test p-value = 0.0003) after incubation with undiluted CupriDyne for 30 minutes. Further incubation with undiluted CupriDyne for 60 minutes reduced viral titers below the limit of detection (< 75 TCID50 per ml).

**Figure 1:**
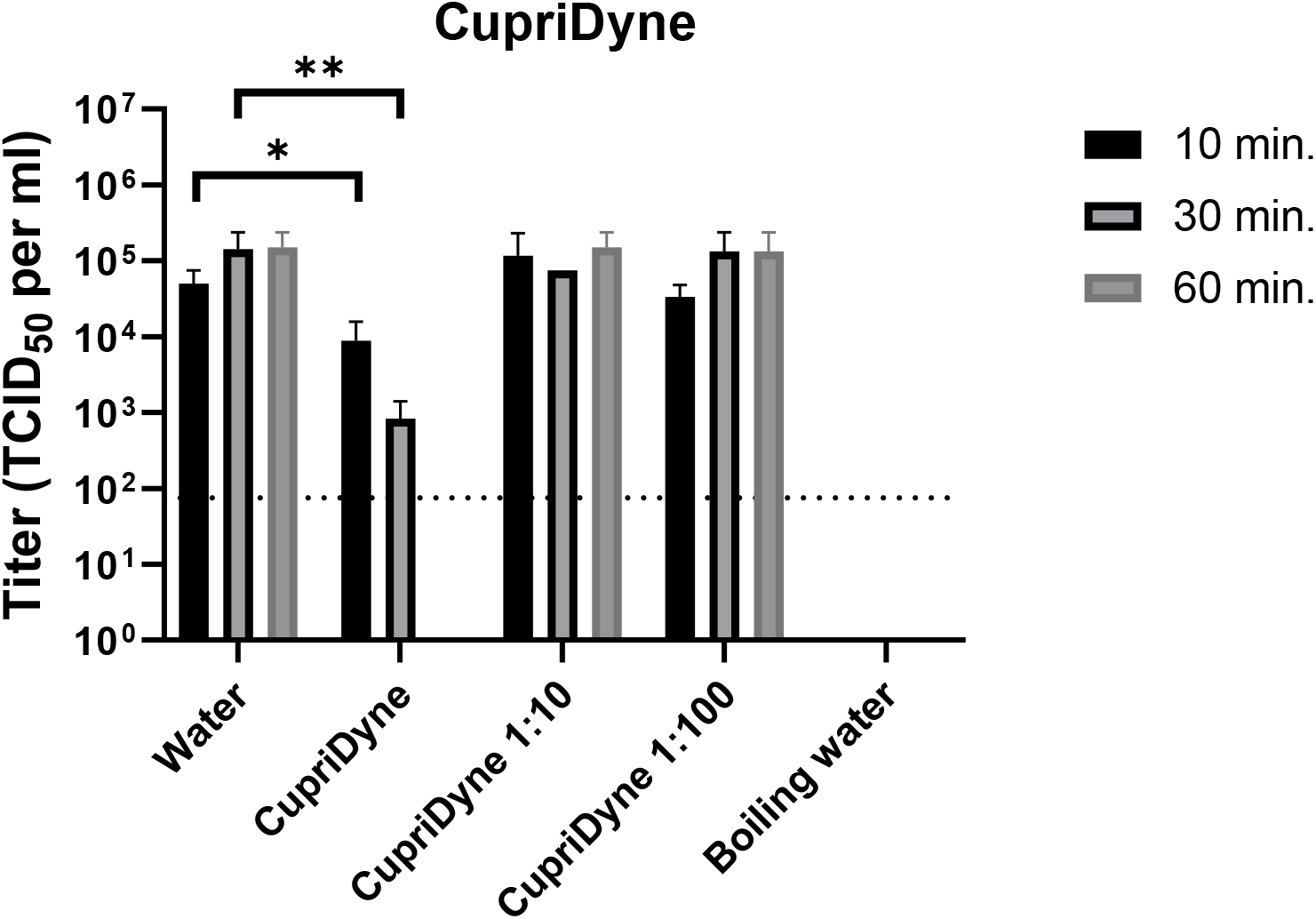
Inactivation of SARS-CoV-2 by CupriDyne. Stocks of SARS-CoV-2 virus were incubated with CupriDyne solutions at 250 ppm (undiluted), 25 ppm (1:10 dilution), or 2.5 ppm (1:100 dilution), room temperature water, or boiling water (>100° C water) for 10, 30, or 60 minutes. Samples were incubated with Vero cell monolayers to determine remaining viable viral particle titers (TCID_50_).

## Discussion

In this study, the efficacy of the CupriDyne iodine complex was assessed for its ability to inactivate SARS-CoV-2. The results clearly show that the CupriDyne iodine complex has viricidal activity against SARS-CoV-2. This effect is both concentration- and time-dependent. The CupriDyne iodine complex may provide an alternative means of addressing and mitigating risks associated with surface contamination and environmental exposure to infective SARS-CoV-2 virus particles.

It is thought that CupriDyne inactivates viruses through the combined effects of iodine and cuprous iodide. The results collected here extend previous work demonstrating antiviral iodine activity against human CoV viruses including SARS-CoV and MERS-CoV to now include SARS-CoV-2 (Eggers *et al*., 2015, Eggers *et al*., 2018). The limited literature available on viral iodine inactivation suggests iodine attacks tyrosine and histidine residues on the viral protein coat resulting in structural changes and loss of infectivity (Gottardi, 1999; Gottardi, 2014). Further knowledge of the mechanism(s) of iodine inactivation may allow for improvements to the viricidal activity of the CupriDyne iodine complex.

Both bleach and alcohol exhibit instability in the environment over time, creating a gap for long-lasting viricidal disinfectants. To determine whether CupriDyne may offer a longer-lasting alternative to bleach and alcohol-based disinfectants, further studies should be conducted to assess CupriDyne’s viricidal efficacy over extended (24 hours and more) exposure times with SARS-CoV-2. Furthermore, additional studies should be conducted to directly compare CupriDyne’s efficacy over time relative to the efficacy of common bleach and alcohol-based solutions. However, initial results clearly show that CupriDyne exhibits viricidal activity against SARS-CoV-2 over 60 minutes.

The CupriDyne iodine complex contains ingredients that are safe for human exposure and which are not known to be associated with poor environmental outcomes. Furthermore, a recent study conducted by a government-certified South Korean laboratory demonstrated that CupriDyne passed all safety, aquatic toxicity, and skin sensitivity tests required by the South Korean government (data not shown). These attributes offer a potential advantage to currently available solutions for environmental control of SARS-CoV-2 such as bleach or alcohol-based products that have downsides for widespread use including skin sensitivity, inhalation risks, and poor environmental outcomes. Further studies should be conducted to assess, at large scale and with good laboratory practice (GLP) standards, any potential negative health or environmental impacts of CupriDyne to verify whether it could represent a safer, more environmentally friendly surface disinfection alternative to conventional solutions.

This study provides strong evidence to the potential suitability of the CupriDyne novel iodine complex for minimizing environmental exposure to the SARS-CoV-2.

## Acknowledgments

We thank Drs. Kenneth Plante (The World Reference Center for Emerging Viruses and Arboviruses, UTMB) and Natalie Thornburg from the CDC for providing the SARS-CoV-2 stock virus. E.K.M was supported by NIH T32 training grant AI060549.

